# Streamline CNV – A Handy Nextflow Pipeline for Chromosomal Copy Number Variation Analysis

**DOI:** 10.64898/2026.10.18.753004

**Authors:** Duanduan Ma, Allen Soberanes, Faith S. Kim, Aditi Shirke, Xiaofeng A. Su

## Abstract

**Background:** Aneuploidy, characterized by imbalanced chromosomal copy number variation (CNV), occurs in both normal somatic tissues and a wide range of diseases. Low pass whole genome sequencing (LP-WGS) to detect CNVs in cells and tissues has been widely used in all biological research. Existing informatics tools for detecting CNV often require computational expertise, often posing challenges for wet-lab biologists. To address this need, we developed Streamline CNV (SLCNV) - a user-friendly and flexible copy number analysis pipeline that allows researchers to perform many copy number-related analyses steps with a single, straightforward command.

**Results:** Streamline CNV is based on Nextflow and is packaged in a Docker image that can be executed using Singularity, making it portable and reproducible across different user environments. The pipeline automatically processes input data, performs sequencing quality control to filter problematic samples, outputs mapping statistics, CNV calling, and generates outputs including intuitive CNV plots, segmentation files (with statistics), annotation of genes on altered chromosomes, evaluation of proportion of chromosomal regions that are non-euploid. Based on user preferences, it can cluster samples solely based on CNV patterns, map the genomic location and density of genes of interest along chromosomes, and visually highlight regions with evident CNVs. In addition, it allows users to select and order chromosomes for display in plots, define sample label color palettes, and choose whether to include sample labels in the clustering visualizations. It also allows users to specify the amount of computing resources to allocate to the pipeline, such as CPU, memory, and runtime. Streamline CNV can be customizable and upgradable. It works with any species and genome assembly and efficiently supports both single-cell and bulk sequencing data with improved performance coding. Jobs can be distributed across cores of a basic Unix workstation, multiple nodes in a high-performance computing cluster using the Slurm scheduler, or cloud environments, as needed.

**Conclusions:** We demonstrated the tool using an *in-house* sequencing dataset of human and mouse cell lines. Our results showed that the Streamline CNV pipeline effectively detected copy number patterns and generated informative summaries. It annotated the findings by identifying affected genes within the copy-number segments and at the break-end regions. It also reported the proportions of altered chromosomal regions impacted by copy number changes for each sample. The clustering plot demonstrated the relationships between samples based on their copy number profiles. Gene density plots aid in interpreting how genes from distinct pathways are associated with CNV pattern alterations. This tool enables researchers, regardless of bioinformatics expertise, to perform comprehensive CNV analyses across a wide range of scenarios. It also facilitates the interpretation of the copy number results and makes it possible for transforming raw CNV data into further exploration of meaningful biological insights.

## Background

Chromosomal copy number variation (CNV), which often causes imbalance in a genome, a phenomenon known as aneuploidy, is a fundamental genomic abnormality occurring in numerous diseases, including over most of the aggressive cancers [1–3] and many genetic diseases which often cause developmental disorders, such as Down syndrome (Trisomy 21) [4], Edwards syndrome (Trisomy 18) [5], Patau syndrome (Trisomy 13) and Turner syndrome (loss of one X chromosome in female or loss of Y chromosome in male), which usually cause severe cellular and physical consequences [6, 7]. CNVs have been found in various chromosomal regions (focal CNVs) in various developmental disorders, such as autism spectrum disorder [8], intellectual disability such as Williams syndrome [9], and attention deficit hyperactivity disorder (ADHD) [10]. Aneuploidy also occurs in normal tissues such as mammalian liver [11] and neurons [12] much less frequently than those diseased tissues [13].

Low-pass whole genome sequencing (LP-WGS) has been widely used for detecting chromosomal CNV in both basic science and clinical settings for cellular and non-cellular DNA samples [14, 15]. A number of computational tools have been developed to facilitate the calling of CNVs based on data from LP-WGS. For example, CNVkit [16], CNVnator [17], GATK gCNV [18], and Control-FREEC [19] are widely used, well-supported, and flexible tools for CNV calling from whole-exome or whole-genome sequencing data. AneuFinder has been used for detecting CNV using single-cell whole genome sequencing (scWGS) [20]. For single-cell(sc) CNV inference from scRNA sequencing, numerous tools such as InferCNV [21], CopyKAT [22], SCEVAN [23], Numbat [24], and CaSpER [25] have been developed and widely used depending on expression with or without allele information (benchmarked in [26, 27]). Single-cell ATAC sequencing has also been used for inferred CNV with CNV callers such as epiAneufinder [28] and copy-scAT [29]. However, while most of these tools are designed to be flexible and accessible to computational biologists, their installation, dependency management, and the execution of a series of commands in the workflows can present a considerable barrier and inconvenience to many experimental biologists without a computational background.

Moreover, a unified solution integrating all critical steps is lacking. Steps ranging from sequencing quality control (QC), filtering problematic samples, sequencing read alignment, CNV detection, segment/gene annotation, summary and statistics, sample clustering based on CNV patterns, to re-coloring plots based on thresholding shallow alterations, visualizing the association between gene density across pathways and CNV patterns, and flexible execution on platforms from basic UNIX computers to high-performance clusters and cloud systems. Existing tools also fall short in offering broad applicability across species, genome assemblies, and sequencing types (from bulk to single cell).

In this study, we introduce **Streamline CNV (SLCNV)**, a CNV analysis pipeline built using the Nextflow framework [30], which seamlessly links all essential steps of a high-quality CNV workflow [13, 31] and additional QC, summary and visualization features that can be run with a single command. To ensure portability and reproducibility, we provide a Docker image containing all dependencies, executable through Singularity [32]. This design makes this pipeline easy to deploy and use: whether on a personal workstation, a high-performance computing cluster, or in the cloud, by any researchers with or without bioinformatics experience.

## Implementation

The earlier in-house CNV pipeline [13] was based on HMMcopy and implemented in Perl and R. It includes basic copy number detection steps (steps 2-4 in the major workflow section) that require computational expertise to execute. We rewrote the original Perl scripts in Python and added essential features for quality control and for converting CNV results into biological insights, as described in the major workflow section. We orchestrated each step using Nextflow [30], enabling the entire workflow to be executed with a single command. We also built a Docker image that incorporates all the scripts and dependencies. Following the manual, the users will be able to install and run the entire flow through Singularity.

SLCNV workflow is described in Figure 1 and further explained in the below steps. The users have an option to distribute the jobs across cores on a basic UNIX workstation, on a single compute node of a computing cluster, or to scatter the jobs across the compute nodes on a computing cluster or cloud using job schedulers like Slurm or equivalents. The parallelization makes it possible to analyze a large number of samples within a short amount of time.

**Figure 1.**
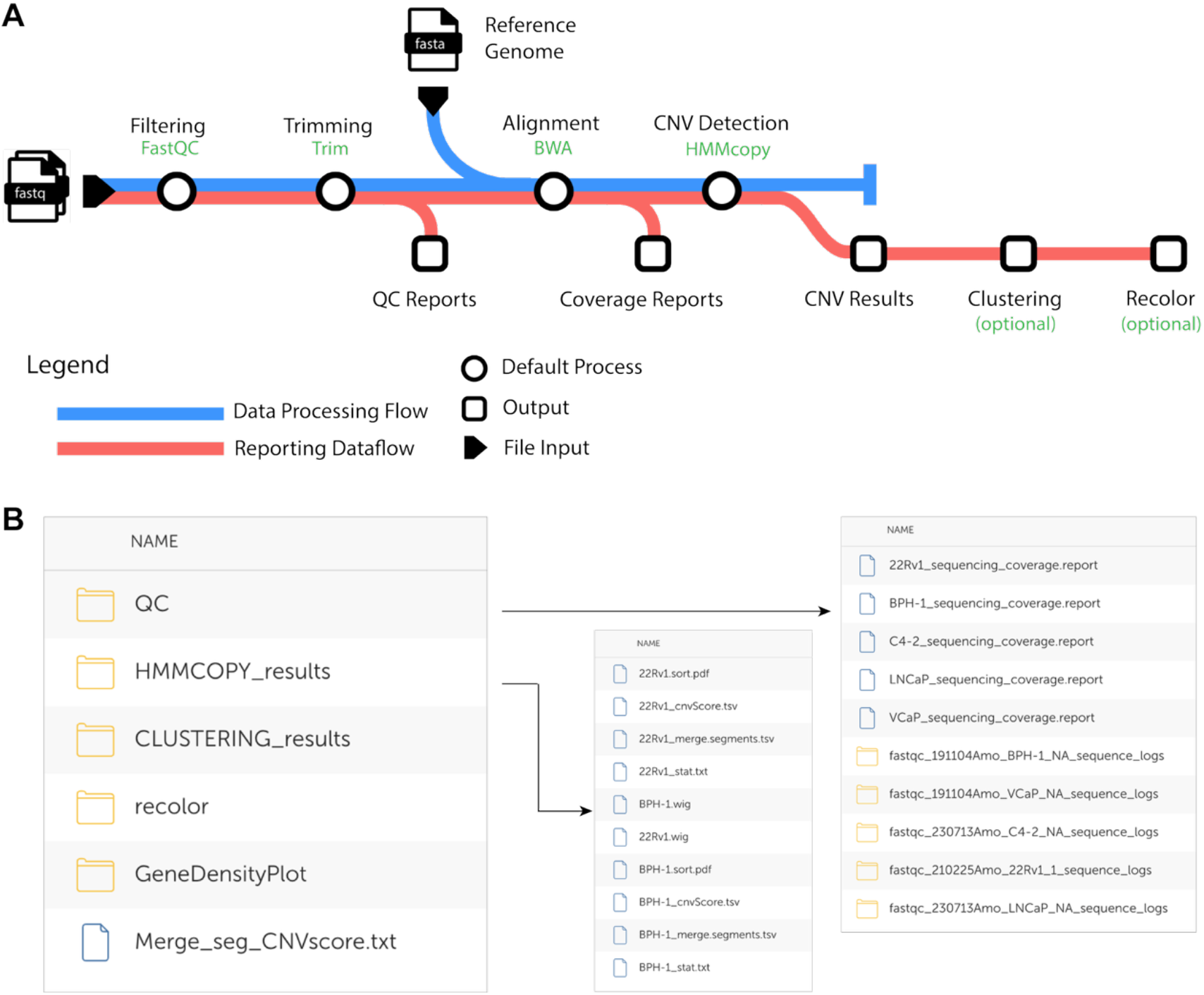
SLCNV pipeline and output delivery. (A) Diagram of SLCNV. Add-on function (i.e. Gene Density visualization) is not included in the main pipeline flow. (B) Layout of an example data output delivery.

## The major workflow

The core workflow of SLCNV consists of five main steps:

1. Quality control of input FASTQ files and removal of problematic samples
2. Trimming of FASTQ sequences
3. Mapping the trimmed reads to the reference genome and generating coverage report
4. Performing CNV analysis using HMMcopy
5. Annotating CNV segments and genes on altered chromosomal regions

Further details for each step are provided below:

### QC and filtering

FastQC was used to assess the quality of the input FASTQ files. Samples with fewer than 50,000 sequencing reads were excluded from downstream analyses [31], as the coverage will be extremely sparse and they usually fail to contain sufficient data to support reliable CNV detection.

### Trimming

The core algorithm underlying SLCNV is HMMcopy, which corrects for mappability and GC content bias in non-overlapping windows of fixed length. To ensure compatibility with the mappability file, typically generated for a specific read length, FASTQ sequences are trimmed to match this length. In SLCNV, all reads are required to be trimmed to 40 bp, ensure the consistency of processing between long and short sequence reads.

### Mapping

The trimmed sequences were aligned to the reference genome using the BWA backtrack algorithm [33]. The resulting SAM alignment files were converted to BAM format using SAMtools [34], followed by sorting and indexing with SAMtools to prepare the data for downstream analysis. Genome-wide coverage after trimming was subsequently reported.

### CNV analyses

CNV analysis was performed using HMMcopy. Additional statistical processing and visualization followed the workflow described in reference [13], with the original Perl scripts re-implemented in Python for improved readability and integration.

### Annotation and summary

Segments with copy number variations were annotated with Ensembl gene IDs corresponding to genes located within each segment. Genes overlapping the breakpoints of segments were also identified and reported. Additional summary analyses were conducted, including calculation of the CNV scores, which represent the proportion of altered chromosomal regions for each chromosome and across the entire genome. These tasks were performed using customized Python scripts.

## Options

Based on user preferences, SLCNV can generate sample clustering plots, gene density plots, highlight regions with significant CNV changes, distribute jobs across multiple compute nodes in a cluster environment, define CPU, memory usage, and runtime, and allow users to specify the reference genome and assembly. Additional functionality is described in the user manual.

Sample Clustering Plot: If users enable the sample clustering option and provide a list of samples of interest, SLCNV will cluster these samples and generate a clustering heatmap. This helps users assess the similarity of input samples based on their CNV patterns. Users have the option to define which chromosomes are displayed, select the sample label color scheme, and decide whether to display sample labels.

### Gene density plots

This is an add-on function of the SLCNV pipeline, which can be run independently from the main processing flow (Figure 1). If users choose to enable gene density plotting and provide a list of genes of interest, SLCNV will visualize each gene as a semi-transparent dot positioned along the chromosomes. This allows users to easily identify regions of high or low gene density and assess their relationship to areas with CNV alterations. To enhance clarity, the figure includes two plots with different transparency levels, providing better visual contrast that facilitating interpretation.

### Recoloring

Segments with CNV changes are visualized using distinct colors called by the default setting of HMMcopy. However, users can enable the recolor option to highlight regions with more pronounced CNV changes. This customization allows for clearer distinction of recurrent CNV regions in the final plots. Caution: performing this function will lose highlights of shallowed gains or losses which could represent small subpopulations in bulk sequencing.

### Job distribution

By default, SLCNV distributes jobs across cores on a local UNIX workstation, a single compute node, or a cloud instance. For users with access to high-performance computing clusters that use the Slurm scheduler, the pipeline supports scattering jobs across multiple compute nodes, enabling greater parallelization and improved processing speed. The users can also define the usage of computing resources including CPU, memory, and run time.

### Species and assembly

SLCNV can analyze CNV changes in any species and genome assembly, based on the user’s choice. By default, SLCNV includes reference data, index files, mappability files, and GC content files for human (GRCh38) and mouse (GRCm39). To analyze other species or assemblies, users must provide their own reference genome, index files, mappability file, and GC content file, following the guidelines provided by HMMcopy utils (https://github.com/shahcompbio/hmmcopy_utils) and the user’s manual.

## Results

Following the SLCNV pipeline shown in Figure 1A, we analyzed 37 human cell lines which were previously sequenced at the MIT BioMicroCenter (BMC) core facility. These cell lines were divided into eight categories: Breast Cancer Line (4 lines), Ewing Sarcoma Cancer Line (4 lines), Glioblastoma Cell Line (8 lines), Normal Human Astrocyte (4 lines), Normal Human Fibroblast (6 lines), Primary Human Mesenchymal Progenitor Cells (3 lines), Primary Prostate Epithelial Cells (2 lines), and Prostate Cancer Cell Lines (6 lines) (See Supplemental Table S1). The data output delivery was shown in Figure 1B as an example.

### QC

QC results are located in the “QC” subfolder within the results directory (Figure 1B). This folder contains the FastQC report and post-trimming sequencing coverage report for each sample. Our current data exhibits the post-trimming coverages ranging from 0.014 to 0.53 for all our samples (Supplemental Table S1). Additionally, warning files are generated if there were any problematic samples that were filtered out. These warning files report the number of reads in the FASTQ files and explain why the problematic samples were removed from downstream analyses.

### CNV results

Most of the CNV results are located in the “HMMcopy_results” subfolder within the results directory. It contains the key outputs of the SLCNV pipeline (Figure 1B). This includes a CNV pattern plot in PDF format and a CNV segment summary table generated by HMMcopy [13, 31] in TSV format for each sample (See examples in Figure 2A and Supplemental Table S2 which provides segment boundaries, CNV states and median values [17,18]). The CNV segment summary table also includes annotations of genes located within aneuploidy segments and at breakpoints (Supplemental Table S2; Note that each sample will generate a segment file which will be listed as an individual tab in Table S2).

**Figure 2.**
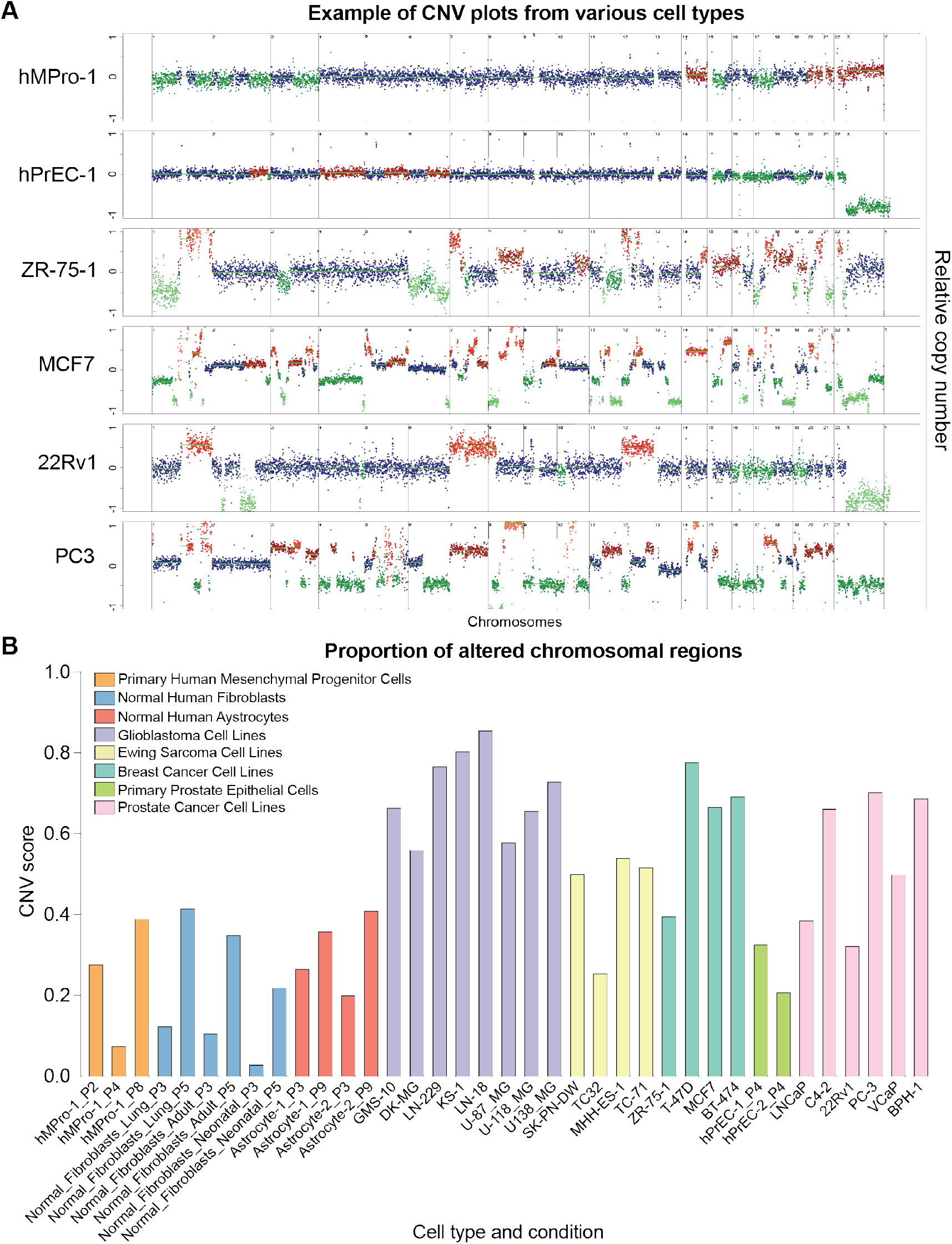
Data plots generated by SLCNV. (A) Examples of CNV plots for the indicated cell lines. All samples tested in this paper is listed in Supplemental Table S1. Part of data were published in [35]. (2) CNV score, defined by proportion of altered chromosomal region, for all the tested samples in this study.

The statistical summary file in text format (.txt) reports metrics including the total number of sequences, number of mapped reads, percentage of mapped sequences, and the variability score (VS) [13]. (Supplemental Table S2; Note that each sample will generate a segment file which will be listed as an individual tab in Table S2).

The CNV score file, in TSV format, indicates the proportion of each altered (either gained or lost) chromosome, as well as the proportion of all altered chromosomal regions as one whole-genome CNV score. Please see the delivery folder example in Figure 1B and the summary of all CNV scores for all the samples (Figure 2B).

The wiggle (WIG) files for each sample are also provided to support visualization and further downstream exploration by the users.

### Clustering

Unsupervised clustering results based on CNV patterns are located in the **“**CLUSTERING_results” subfolder within the results directory. As shown in our example data (Figure 3), samples from the same cell or cancer type with similar CNV scores tend to cluster together. Interestingly, our results showed that cells from the same cell line often exhibit distinct subclusters due to differences in passage number, which is correlated with increased CNVs (Figure 2B) associated with increased passages *in vitro*. We’ve also observed that prostate cancer cell lines tend to have relatively lower CNV scores and cluster together consistent with previous cancer case analysis [2]. Moreover, the cells from the same cancer type may exhibit unique CNV patterns, highlighting both the aneuploidy changes that occur during cancer evolution and the inherent variability among individual cancer cells.

**Figure 3.**
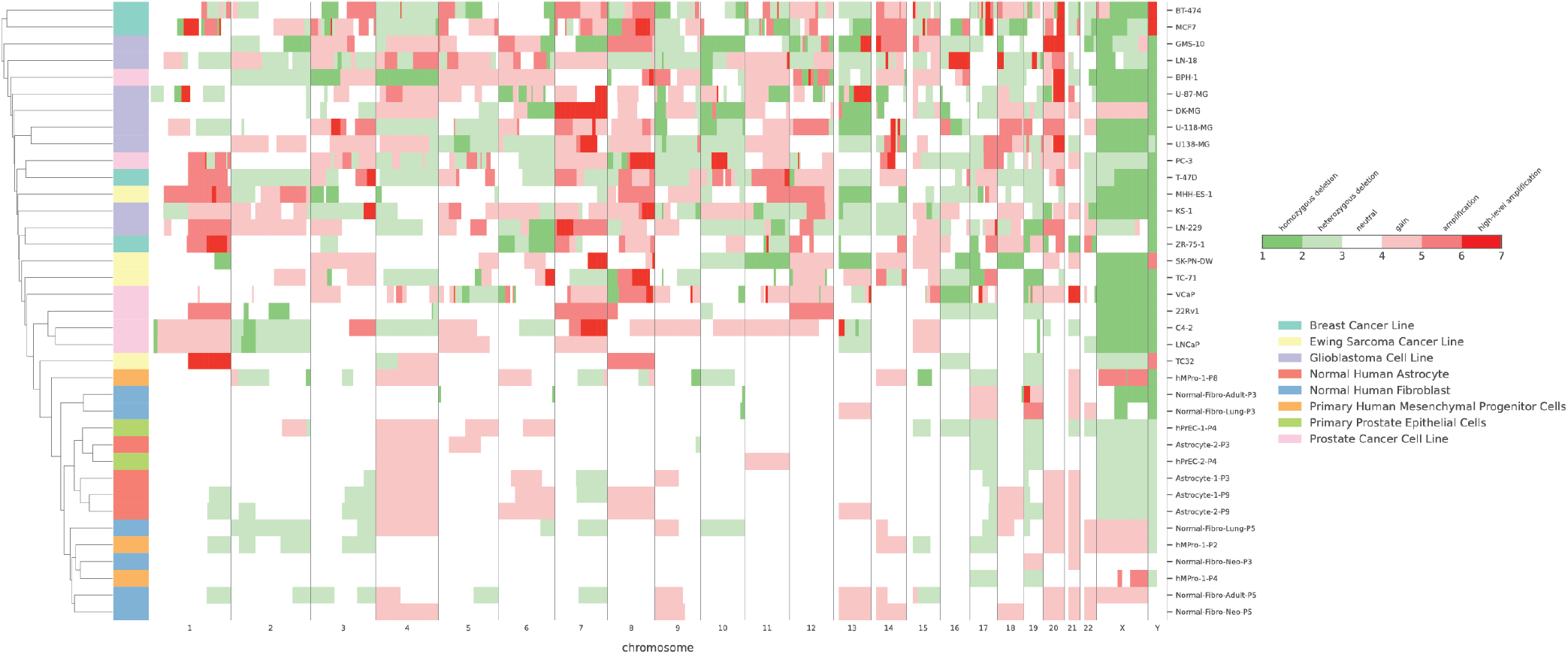
Un-supervised clustering based on CNV patterns for all the tested samples in this study.

### Recolor

The recolored plots are located in the “recolor” subfolder within the results directory. More pronounced CNV patterns are highlighted to draw additional attention (Figure 4).

**Figure 4.**
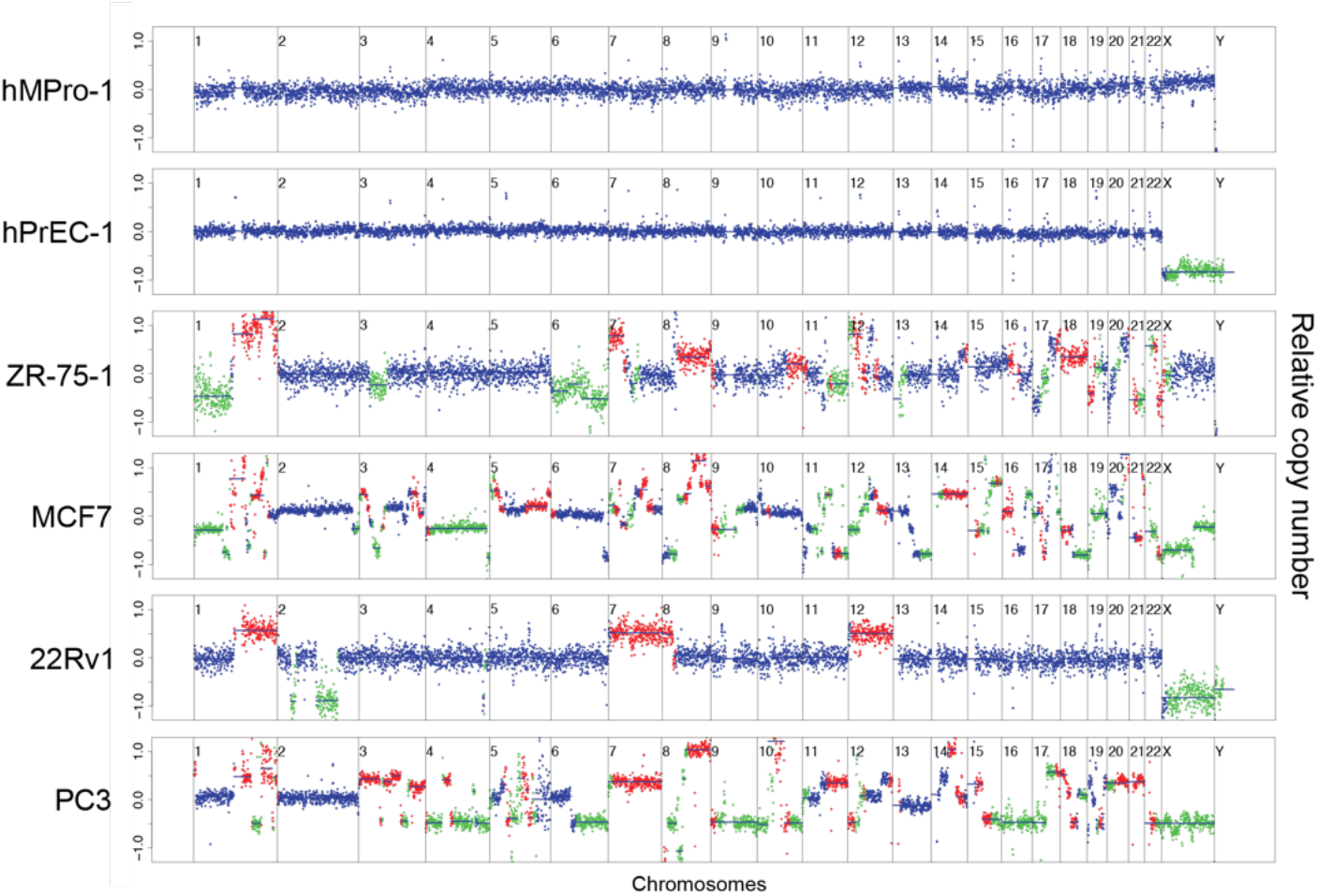
Re-coloring CNV plots in the examples shown in Figure 2A.

### Gene density visualization

This is an add-on function independent from using FASTQ files to determine CNV, that helps relate CNV patterns to their underlying biological mechanisms. For example, given a list of key genes in a pathway (e.g., cell cycle regulation), visualizing their density along each chromosome and comparing it with the locations of aneuploidy segments can help elucidate the involvement of relevant pathways underlying CNV changes.

## Conclusions

SLCNV is a flexible and user-friendly pipeline designed to support both biological and clinical research communities. It delivers reliable CNV detection, annotation, visualization, and summary results. By integrating quality control, CNV detection, annotation, summarization, clustering and visualization, SLCNV enables not only the identification of CNV changes in single cells and bulk samples, but also the visualization of longitudinal CNV dynamics during development, disease progression, cancer clonal evolution, and cell line passage. Furthermore, it facilitates the understanding of underlying biological mechanisms by identifying affected genes and pathways. SLCNV serves as a practical, all-in-one tool that brings together key analytical components, accessible to users regardless of their bioinformatics or computational background.

## Availability and requirements

### Project name

SLCNV

### Project home page

https://github.com/allensulab/StreamlineCNV

### Operating system(s)

UNIX

### Programming language

Nextflow (Groovy), Python, R

### Other requirements

Docker, Singularity

### License

NA

### Any restrictions to use by non-academics

NA

## Supporting information

User_Manual

## List of abbreviations

SLCNV: Streamline
CNV CNV: Copy Number Variation

## Declarations

### Ethics approval and consent to participate

Not applicable -- All cells and samples used in this study are publicly available or have been published previously; they were used under material transfer and use agreements with vendors from which they were commercially purchased for research purposes. There is no requirement for ethics approval or informed consent. Please refer: https://www.biomedcentral.com/getpublished/editorial-policies#ethics+and+consent)

### Consent for publication

Not applicable. Please refer the guidelines from BMC bioinformatics at https://bmcbioinformatics.biomedcentral.com/submission-guidelines/preparing-your-manuscript/software-article. Look for the section named “Consent for publication”).

### Availability of data and materials

The datasets used during the current study are available from the corresponding author on reasonable request. The processed (ready-to-use) data are available in Sequence Read Archive (SRA) (BioProject ID: PRJNA1434537).

### Authors’ contributions

**D.M**. designed and developed SLCNV, and drafted the manuscript and user manual. **A.S.O**. created the Docker image, implemented Singularity applications, and edited the user manual. **F.S.K**. applied and tested SLCNV across multiple Unix workstations, computing clusters, and cloud environments and also provided valuable feedback. **A.S**. further develop add-on function. **X.A.S**. initiated this project, and conceived critical ideas underlying SLCNV, co-wrote the manuscript, and provided funding support.

## Acknowledgements

This work has been supported in part by the Koch Institute Support (core) Grant P30-CA014051 from the National Cancer Institute (NCI). We thank the Koch Institute’s Robert A. Swanson (1969) Biotechnology Center for technical support, specifically Genomics and Bioinformatics core facilities. We thank Dr. Amon lab’s support for the postdoctoral work of X.A.S. who was a previous Jane Coffin Childs postdoctoral fellow. X.A.S. are partially supported by Center for Prostate Disease Research. X.A.S. (partially) and A.S. are supported by a Murtha Cancer Center Research Program – PROMETHEUS funding. A part of this work and F.K. are supported by an Idea Development Award (IDA) HT9425-25-1-0255 awarded to X.A.S. from Congressionally Directed Medical Research Programs (CDMRP) Prostate Cancer Research Program (PCRP).

## References

1. Beroukhim R, Mermel CH, Porter D, Wei G, Raychaudhuri S, Donovan J, Barretina J, Boehm JS, Dobson J, Urashima M et al: The landscape of somatic copy-number alteration across human cancers. Nature 2010, 463(7283):899–905.

2. Taylor AM, Shih J, Ha G, Gao GF, Zhang X, Berger AC, Schumacher SE, Wang C, Hu H, Liu J et al: Genomic and Functional Approaches to Understanding Cancer Aneuploidy. Cancer Cell 2018, 33(4):676–689 e673.

3. Weaver BA, Cleveland DW: Does aneuploidy cause cancer? Curr Opin Cell Biol 2006, 18(6):658–667.

4. Antonarakis SE, Skotko BG, Rafii MS, Strydom A, Pape SE, Bianchi DW, Sherman SL, Reeves RH: Down syndrome. Nat Rev Dis Primers 2020, 6(1):9.

5. Cereda A, Carey JC: The trisomy 18 syndrome. Orphanet J Rare Dis 2012, 7:81.

6. Torres EM: Consequences of gaining an extra chromosome. Chromosome Res 2023, 31(3):24.

7. Santaguida S, Amon A: Short- and long-term effects of chromosome mis-segregation and aneuploidy. Nat Rev Mol Cell Biol 2015, 16(8):473–485.

8. Vicari S, Napoli E, Cordeddu V, Menghini D, Alesi V, Loddo S, Novelli A, Tartaglia M: Copy number variants in autism spectrum disorders. Prog Neuropsychopharmacol Biol Psychiatry 2019, 92:421–427.

9. Merla G, Brunetti-Pierri N, Micale L, Fusco C: Copy number variants at Williams-Beuren syndrome 7q11.23 region. Hum Genet 2010, 128(1):3–26.

10. Martin J, Hosking G, Wadon M, Agha SS, Langley K, Rees E, Owen MJ, O’Donovan M, Kirov G, Thapar A: A brief report: de novo copy number variants in children with attention deficit hyperactivity disorder. Transl Psychiatry 2020, 10(1):135.

11. Tremmel R, Klein K, Battke F, Fehr S, Winter S, Scheurenbrand T, Schaeffeler E, Biskup S, Schwab M, Zanger UM: Copy number variation profiling in pharmacogenes using panel-based exome resequencing and correlation to human liver expression. Hum Genet 2020, 139(2):137–149.

12. Wade L: Mercury pollution. Gold’s dark side. Science 2013, 341(6153):1448–1449.

13. Knouse KA, Wu J, Whittaker CA, Amon A: Single cell sequencing reveals low levels of aneuploidy across mammalian tissues. Proc Natl Acad Sci U S A 2014, 111(37):13409–13414.

14. Mazzonetto PC, Villela D, da Costa SS, Krepischi ACV, Milanezi F, Migliavacca MP, Pierry PM, Bonaldi A, Almeida LGD, De Souza CA et al: Low-pass whole genome sequencing is a reliable and cost-effective approach for copy number variant analysis in the clinical setting. Ann Hum Genet 2024, 88(2):113–125.

15. Wang N, Tao ZY, Wu T, Wang J, Wang W, Shi H, Liu XS: Benchmarking copy number variation detection with low-coverage whole-genome sequencing. Brief Bioinform 2025, 26(5).

16. Talevich E, Shain AH, Botton T, Bastian BC: CNVkit: Genome-Wide Copy Number Detection and Visualization from Targeted DNA Sequencing. PLoS Comput Biol 2016, 12(4):e1004873.

17. Abyzov A, Urban AE, Snyder M, Gerstein M: CNVnator: an approach to discover, genotype, and characterize typical and atypical CNVs from family and population genome sequencing. Genome Res 2011, 21(6):974–984.

18. Babadi M, Fu JM, Lee SK, Smirnov AN, Gauthier LD, Walker M, Benjamin DI, Zhao X, Karczewski KJ, Wong I et al: GATK-gCNV enables the discovery of rare copy number variants from exome sequencing data. Nat Genet 2023, 55(9):1589–1597.

19. Boeva V, Popova T, Bleakley K, Chiche P, Cappo J, Schleiermacher G, Janoueix-Lerosey I, Delattre O, Barillot E: Control-FREEC: a tool for assessing copy number and allelic content using next-generation sequencing data. Bioinformatics 2012, 28(3):423–425.

20. Bakker B, Taudt A, Belderbos ME, Porubsky D, Spierings DC, de Jong TV, Halsema N, Kazemier HG, Hoekstra-Wakker K, Bradley A et al: Single-cell sequencing reveals karyotype heterogeneity in murine and human malignancies. Genome Biol 2016, 17(1):115.

21. Tirosh I, Izar B, Prakadan SM, Wadsworth MH, 2nd, Treacy D, Trombetta JJ, Rotem A, Rodman C, Lian C, Murphy G et al: Dissecting the multicellular ecosystem of metastatic melanoma by single-cell RNA-seq. Science 2016, 352(6282):189–196.

22. Gao R, Bai S, Henderson YC, Lin Y, Schalck A, Yan Y, Kumar T, Hu M, Sei E, Davis A et al: Delineating copy number and clonal substructure in human tumors from single-cell transcriptomes. Nat Biotechnol 2021, 39(5):599–608.

23. De Falco A, Caruso F, Su XD, Iavarone A, Ceccarelli M: A variational algorithm to detect the clonal copy number substructure of tumors from scRNA-seq data. Nat Commun 2023, 14(1):1074.

24. Gao T, Soldatov R, Sarkar H, Kurkiewicz A, Biederstedt E, Loh PR, Kharchenko PV: Haplotype-aware analysis of somatic copy number variations from single-cell transcriptomes. Nat Biotechnol 2023, 41(3):417–426.

25. Serin Harmanci A, Harmanci AO, Zhou X: CaSpER identifies and visualizes CNV events by integrative analysis of single-cell or bulk RNA-sequencing data. Nat Commun 2020, 11(1):89.

26. Schmid KT, Symeonidi A, Hlushchenko D, Richter ML, Tijhuis AE, Foijer F, Colome-Tatche M: Benchmarking scRNA-seq copy number variation callers. Nat Commun 2025, 16(1):8777.

27. Song M, Ma S, Wang G, Wang Y, Yang Z, Xie B, Guo T, Huang X, Zhang L: Benchmarking copy number aberrations inference tools using single-cell multi-omics datasets. Brief Bioinform 2025, 26(2).

28. Ramakrishnan A, Symeonidi A, Hanel P, Schmid KT, Richter ML, Schubert M, Colome-Tatche M: epiAneufinder identifies copy number alterations from single-cell ATAC-seq data. Nat Commun 2023, 14(1):5846.

29. Nikolic A, Singhal D, Ellestad K, Johnston M, Shen Y, Gillmor A, Morrissy S, Cairncross JG, Jones S, Lupien M et al: Copy-scAT: Deconvoluting single-cell chromatin accessibility of genetic subclones in cancer. Sci Adv 2021, 7(42):eabg6045.

30. Di Tommaso P, Chatzou M, Floden EW, Barja PP, Palumbo E, Notredame C: Nextflow enables reproducible computational workflows. Nat Biotechnol 2017, 35(4):316–319.

31. HMMcopy: Copy number prediction with correction for GC and mappability bias for HTS data [https://bioconductor.org/packages/HMMcopy]

32. Kurtzer GM, Sochat V, Bauer MW: Singularity: Scientific containers for mobility of compute. PLoS One 2017, 12(5):e0177459.

33. Li H, Durbin R: Fast and accurate short read alignment with Burrows-Wheeler transform. Bioinformatics 2009, 25(14):1754–1760.

34. Danecek P, Bonfield JK, Liddle J, Marshall J, Ohan V, Pollard MO, Whitwham A, Keane T, McCarthy SA, Davies RM et al: Twelve years of SAMtools and BCFtools. Gigascience 2021, 10(2).

35. Su XA, Ma D, Parsons JV, Replogle JM, Amatruda JF, Whittaker CA, Stegmaier K, Amon A: RAD21 is a driver of chromosome 8 gain in Ewing sarcoma to mitigate replication stress. Genes Dev 2021, 35(7-8):556–572.

