## Supplementary material for "Streamline CNV – A Handy Nextflow Pipeline for Chromosomal Copy Number Variation Analysis": User_Manual

**Streamline CNV (SLCNV) User Manual**

### Installation:

All the pipeline tools are packaged into an image which adheres to the Open Container Initiative (OCI) specification. The OCI container image, run scripts, and Nextflow configuration files are hosted on a GitHub repository, https://github.com/allensulab/StreamlineCNV. The pipeline can be executed using one of three provided run scripts, each designed for a different computing environment:

1. `run_local.sh`: for execution on a local machine
2. `run_slurm.sh`: for execution on a high-performance computing (HPC) cluster utilizing the Slurm job scheduler
3. `run_singularity_no_nextflow.sh`: for execution on an HPC system with Singularity available but without Nextflow

This pipeline requires Singularity, and ideally, Nextflow.

Install Singularity by following the official installation instructions. Apptainer, the community-maintained successor to Singularity, may also be used. Likewise, install Nextflow by following the official installation instructions. If using an HPC system, it is recommended to first verify whether these tools are already installed. Please contact your system administrators for more information.

Once Singularity and Nextflow have been installed, clone the GitHub repository to your system:

`git clone https://github.com/allensulab/StreamlineCNV && cd StreamlineCNV`

### Environment Variables

Before running the pipeline, the following environment variables must be defined. This setup enables the pipeline to locate the input data. The example shell scripts are provided for this purpose.

| Variable | Description |
|---|---|
| DATA_DIR | Path to directory containing genome reference files. This directory is mounted at '/data' inside the Singularity container. Defaults to `data/`. |
| FASTQ_DATA | Path to directory containing FASTQ. Defaults to `data/fastq_files`. |
| OUTDIR | Output directory for pipeline results. Defaults to `results`. |
| GENE_LIST | (Optional) Path to the gene list file used for plotting gene density. Defaults to `data/GeneListExample`. |
| SAMPLE_INFO | Path to the sample information file used for recoloring samples. Defaults to `data/SampleInfo`. |
| CLUSTERING_LABEL | Path to the file containing sample IDs and their associated labels (e.g. cell types) for clustering analyses. Defaults to `data/ClusteringLabel`. |
| QUEUE | (For Slurm users only) Name of the Slurm partition to which the job will be submitted. Defaults to your cluster's default partition. |

| Variable | Description |
|---|---|
| CONTAINER_PROFILE | Containerization software used by pipeline. Defaults to `singularity`. |
| EXTRA_MOUNT | (Optional) Additional mount point to provide to the Singularity container. Defaults to ``. |

**File preparation:**

1. Either move the genome reference files and FASTQ data into the appropriate subdirectories within the `data` folder of the cloned repository, or change the corresponding environment variables to reflect their actual locations on your system.
2. Ensure FASTQ files are named using the following format: `RunName_SampleName_ReadDirection_sequence.fastq`.
    a. Examples:
        i. 180103AmoA_D18-124_NA_sequence.fastq
        ii. 180103AmoA_D18-124_1_sequence.fastq,
        iii. 180103AmoA_D18-124_F_sequence.fastq
    b. The FASTQ file directory should contain **only FASTQ files** and no other file types.
    c. SampleName must be unique per each FASTQ file.
    d. It is recommended to use standard characters and numbers for RunName and SampleName.
        i. Characters such as "_", "&", and "$" may result in downstream errors.
3. All the references and necessary files for the human GRCh38 assembly and mouse GRCm39 assembly have already been prepared and available HERE for download. However, if working with a different species or using an alternative assembly of human and mice genome, users will need to download the appropriate reference genome from either NCBI or Ensembl. Next, please follow the Burrows-Wheeler Aligner (BWA) manual and samtools manual to index the reference FASTA file, and refer to the HMMCopy instructions to generate the genome reference mappability file as well as the genome reference GC content file. In addition, minor tweaks of `run_hmmcopy*.r` files in the github repository are required for species other than human and mice. If changes were made in this case, the Docker image must be rebuilt to include the new scripts by following the instructions in the README file under "container/" directory.
4. Gene list file: If `--plotGeneDensity` is set to `true`, a plain text file containing the list of genes to be plotted must be provided.
    a. The file should contain one gene name per line in a single column.
    b. Please refer to the example file `GeneListExample` included in the repository downloaded from GitHub.
5. SampleInfo file: If `--recolor` is set to `true`, a plain text file containing sample information must be provided.
    a. The file should contain three columns with no header.
        i. First column: Assembly name (case-sensitive) matching the assembly section of the reference file name (e.g., GRCh38)
        ii. Second column: Sample name matching the SampleName section in the FASTQ file name (e.g., D18-124)
        iii. Third column: Gender, with values F or M
    b. Please refer to the example file SampleInfo included in the files downloaded from GitHub.
6. ClusteringLabel file: If `--clustering` is set to `true`, a plain text file containing clustering label information must be provided.
    a. The file should contain two columns with a header: `sample` and `label`.
        i. First column: Sample name matching the SampleName section in the FASTQ file name
        ii. Second column: Annotation for each sample such as tissue type or cell name

b. Please refer to the example file `ClusteringLabel` included in the files downloaded from GitHub.

### Running the pipeline

#### Run on a local workstation

Ensure that Singularity/Apptainer and Nextflow are installed. Open the `run_local.sh` file and change the environment variables to reflect the location of the files on your computer. Run `./run_local.sh`.

Pipeline results will be written to the directory specified in `OUTDIR`. By default, this is a folder named `results` in your current working directory.

#### Run on an HPC system with Slurm

Ensure that Singularity/Apptainer and Nextflow are installed or loaded into your environment. Open the `run_slurm.sh` file and change the environment variables to reflect the location of the files on your server.

If the files are located on a storage array that your server mounts using NFS, set the 'EXTRA_MOUNT' option to the path where the NFS mount starts. For example, if your server mounts storage arrays at `/net/` and your data are stored there, set `EXTRA_MOUNT` to `/net:/net` so that these files are accessible inside the Singularity image at the same path as on your server.

Submit the script to Slurm by running `sbatch run_slurm.sh`.

Pipeline results will be written to the directory specified in `OUTDIR`. By default, this is a folder named `results` in your current working directory.

#### Run without installing Nextflow, only Singularity

If for some reason you are able to install Singularity/Apptainer, but not Nextflow, this pipeline can still be executed. However, jobs cannot be distributed across multiple compute nodes, limiting functionality on HPC environments.

Open the `run_singularity_no_nextflow.sh` file and change the environment variables to reflect the location of the files on your server.

Run `./run_singularity_no_nextflow.sh`.

Alternatively, submit the script to Slurm by running `sbatch run_singularity_no_nextflow.sh`. If you do this, ensure that the script includes commands to load Singularity into your HPC environment.

#### Pipeline arguments and options:

Required arguments

- -profile: Specifies the Nextflow profile(s) to use. Multiple profiles can be used by passing each profile with a comma-separated list.
    - -profile singularity: Run using the Singularity/Apptainer container which contains all pipeline tools and scripts.
    - -profile slurm: Scatter jobs across compute nodes in an HPC system.
    - -profile local: Run on local workstation without scattering to compute nodes.
    - -profile singularity_no_nextflow: Run on local workstation but only when using the Nextflow program installed INSIDE the container.

- --data_dir: Path to the directory containing reference files.
- --fastq: Path to the directory containing user-inputted FASTQ files. This directory must contain only FASTQ files.
- --species: Species to be analyzed. Value must exactly match the name of the reference FASTA file (case-sensitive). For example, if the reference genome is named Homo_sapiens.GRCh38.dna.primary_assembly.fa, then set --species ‘Homo_sapiens’.
- --assembly: Genome assembly. Value must exactly match the name of the reference FASTA file (case-sensitive). For example, if the reference genome is named Homo_sapiens.GRCh38.dna.primary_assembly.fa, then set --assembly ‘GRCh38’.
- --outdir: Path to the output directory for pipeline results.

Optional features

- --plotGeneDensity: Whether to generate gene density plots. Accepts `true` or `false` (default is `false`).
- --geneList: Path to the file containing gene names for gene density plots. Applies only if --plotGeneDensity option is set to `true` (see Gene list file in the File preparation section).
- --recolor: Whether to recolor CNV plots to highlight segments with significant copy number changes. All shallow changes will be recolored as euploid (no change). Accepts `true` or `false` (default is `false`).
- --sampleInfo: Path to the file containing sample information for recoloring. Applies only if --recolor option is set to `true` (see SampleInfo file in the File preparation section). If the sample ID is not included in the input fastq files, or if the fastq file is filtered out due to too few sequencing reads, the nonexistent sample will not be processed, and a warning message will be output in the `recolor` sub-folder inside the result output folder.
- --clustering: Whether to perform clustering analysis based on CNV patterns. Accepts `true` or `false` (default is `true`).
- --clusteringLabel: Path to the file specifying samples and annotations for clustering. Applies only if --clustering option is set to `true` (see ClusteringLabel file in the File preparation section).
- --clustering_memory: Specifies the amount of RAM allocated for clustering (default is 16 GB). Users can customize this by setting a different value, e.g., --clustering_memory ‘64.GB’
- --clustering_cpus: Specifies the number of CPUs allocated for clustering (default is 4). Users can customize this by setting a different value, e.g., --clustering_cpus 8
- --clustering_time: Specifies the maximum runtime for clustering in hours (default is 10). Users can customize this by setting a different value, e.g., --clustering_time ‘20.h’
- --dropChr: Specifies chromosomes to exclude from the output clustering heatmap and CNV plots (default is none ‘ ‘). The listed chromosomes must be space-separated, e.g., --dropChr ‘X Y 19’
- --tissue_color_palette: Matplotlib color palette used for sample labels in the clustering plot (defaults to Set3).
- --no_sampleLabel: Whether to remove sample names along the y-axis of the output clustering heatmap. Accepts `true` or `false` (default is `false`).
- --queue: Name of the Slurm partition for job submission. Applies only when -profile slurm is used.
- --extra_mount: Name of the optional additional mount point.

- --resume: Whether to resume the pipeline from where the previous run was terminated. Without this option, the pipeline starts from the beginning.

**Pipeline outputs:**

The output result folder contains the following sub folders:

QC:

The QC subfolder contains the FastQC results and sequencing coverage information of each sample. If there is a FASTQ file with less than 50,000 sequences, this FASTQ file will be filtered away from downstream analyses. A warning file will be output to the QC folder with the suffix of the samples IDs corresponding to the samples that are filtered away.

HMMCOPY_results:

The HMMCOPY_results subfolder contains the major CNV results.

- The pdf files are the plots for CNV pattern visualization.
- The merge.segments.tsv files contain the detailed information of the CNV segments
- The stat.txt files contain the statistical results of each sample such as the number of total reads, the number of mapped reads, the percentage of mapped sequences, and VS scores that decide whether the CNV results are reliable.
- The cnvScore.tsv files provide information of the proportion of aneuploidy on each chromosome and across the entire genome. Information from all samples will be collected in a summary `Merge_seg_CNVscore.txt` file found in the `results` directory.
- The wig files provide detailed coverage information across the entire genome for further downstream analyses and visualization.

CLUSTERING_results:

If --clustering option is set to `true`, a file named clustering.pdf is generated, displaying a heatmap of the samples defined in the ClusteringLabel file.

recolor:

The pdf files contain the re-colored CNV plots for the samples defined in the SampleInfo file and if --recolor option is set to `true`. If there are sample IDs defined in the SampleInfo files that do not exist or were filtered away, they are skipped during the recolor process and output in warning.txt files.

GeneDensityPlot:

If –plotGeneDensity option is set to `true`, the .jpg files display the gene density information across the chromosomes for the genes defined in the GeneListExample file.
